# Global spread and evolutionary history of HCV subtype 3a

**DOI:** 10.1101/2021.02.03.429581

**Authors:** Shang-kuan lin, Nicola De Maio, Vincent Pedergnana, Chieh-Hsi Wu, Julien Thézé, Ellie Barnes, M. Azim Ansari

## Abstract

Studies have shown that HCV subtype 3a had likely been circulating in South Asia before its global spread. However, the time and route of this dissemination remain unclear. For the first time, we generated host and virus genome-wide data for more than 500 patients infected with HCV subtype 3a from the UK, North America, Australia and New Zealand. We used the host genomic data to infer the ancestry of the patients and used this information to investigate the epidemic history of HCV subtype 3a. We observed that viruses from hosts of South Asian ancestry clustered together near the root of the tree, irrespective of the sampling country and that they were more diverse than viruses from other host ancestries. We also inferred that three independent transmission events resulted in the spread of the virus from South Asia to the UK, North America and the Australian continent. This initial spread happened during or soon after the end of the second world war. This was followed by an exponential growth in the effective population size of HCV subtype 3a worldwide and many independent transmissions between the UK, North America and Australian continent. Using both host and virus genomic information can be highly informative in studying the virus epidemic history especially in the context of chronic infections.

## Introduction

It is estimated that worldwide 71 million people are infected with Hepatitis C virus (HCV), many in the developing world (1). Following the acute phase of infection, the majority of the infected individuals enter a chronic asymptomatic phase of infection that can last decades. Chronic HCV infection is one of the leading causes of liver cirrhosis and hepatocellular carcinoma (HCC) worldwide (2). While there are currently no vaccines for HCV, the recently developed direct-acting antivirals have resulted in significant improvement in the safety and efficacy of treatment regimens for HCV infection.

HCV is highly diverse and is currently classified into eight major genotypes (denoted by numbers 1 to 8), each of which has been divided into many subtypes (denoted by lower case letters e.g. 1a, 1b, etc.). The clinical outcomes of chronic HCV infection are heavily influenced by viral genetics. For example, it has been shown that HCV genotype 3 is associated with higher risk of developing HCC (3–6) and with lower rates of treatment success for direct-acting antivirals (7,8). Furthermore, amino acid variation in the NS5A protein is associated with viral load (9,10) and amino acid variation in the core protein is associated with the development of HCC (11).

The various genotypes and subtypes of HCV are associated with distinct epidemiological and geographical patterns of distribution. Many HCV genotypes have limited geographical distribution. For example, genotype 4 is found mostly in the Middle East and Central Africa, genotype 5 in Southern Africa, and genotype 6 in East and South-East Asia (12). On the other hand, a few HCV subtypes including subtypes 1a, 1b and 3a are globally distributed and have caused most of the HCV infections worldwide (12). Phylogenetic studies have found regions of potential origin for some of these globally distributed subtypes. For example, HCV genotype 1 isolates sampled from Central and West Africa have much higher genetic diversity than those sampled from the other parts of the world (13,14) indicating a long-term endemicity in the region, followed by the global spread of subtype 1a. For HCV genotype 3, studies have found the virus to be highly diverse in South Asia and South-East Asia, indicating the origin of the global epidemic of subtype 3a (15,16). Aside from the investigations of the origins of different genotypes, various studies have also inferred their evolutionary and epidemiological history (17–23). Most of these studies have focused on specific genomic regions (such as Core, E1/E2, NS5A/NS5B) and/or the epidemiological history on restricted geographic regions and have used limited numbers of samples.

In this study, we investigated epidemiological history and global spread of HCV subtype 3a (HCV-3a) using virus whole genomes collected from 507 patients in the United Kingdom, Canada, USA, Australia and New Zealand (BOSON cohort (24)),

Additionally, we generated host genetic data (genome-wide genotyping) for patients in this cohort. This additional source of information allowed us to account for confounding caused by differences between virus sampling locations and the locations of host infection which can be different in the context of chronic infections such as HCV. Combining the host and virus genomic information can improve the genetic epidemiology of chronic viral infections and reduce confounding errors due to migration history of individuals.

## Result

### Molecular clock signal of HCV-3a sequences

The 507 HCV-3a sequences from the BOSON clinical trial were collected during 2013-2014 period (24). Due to the short collection interval, these sequences lack temporal signal of measurable evolution (Supplementary Figure 1). To address this, we added 48 previously published HCV-3a whole genomes, the earliest of which was collected in 1991 in Canada. To explore the temporal signal of the updated dataset, we estimated a maximum likelihood tree from whole genomes and calculated the correlation between root-to-tip genetic distance and sampling dates of the sequences (Supplementary Figure 1). The results showed sufficient molecular clock signal with an estimated substitution rate of 2.13×10^−3^ (95% CI: 1.61×10^−3^-2.64×10^−3^) per site per year, which is larger than previous estimate for HCV-3a (1.65×10^−3^ substitutions per site per year, with 95% HPD: 1.19×10^−3^ −2.14×10^−3^ (19)). However, correlation between root-to-tip distance and sampling dates assumes a strict clock model. A more sophisticated molecular clock estimation was conducted using a Bayesian approach in BEAST (25) assuming a relaxed clock model. The estimated substitution rate was 1.69×10^−3^ (95% HPD: 1.41×10^−3^-1.96×10^−3^) substitutions per site per year.

### Enrichment of South Asian ancestry individuals among HCV-3a infected patients

The principal components (PCs) of host genome-wide genotyping data for the BOSON cohort were projected onto the genetic PCs of the 1000 Genomes project (26). These projections were then used to validate and adjust the self-reported host ancestry of BOSON cohort patients (Figure 1, Supplementary Figure 2, see methods) and allowed us to distinguish between South Asian and East Asian host ancestry (both were self-reported as Asian). For BOSON hosts with HCV-3a infection where genetic data were not available, host ancestries were designated as their self-reported ancestries (19 individuals). The host ancestries corresponding to HCV-3a sequences downloaded from the public repositories (where we do not have access to host genetic data) were designated as the majority ethnic group of the country where the samples were collected. We observed that majority of the patients had white European ancestry (N=433, 80%) and South Asians were the second largest group (N=69, 13%). The other host ancestries present in this dataset were of Admixed American (N=12), East Asian (N=6), African (N=4) and other ethnic groups (N=16).

**Figure 1:**
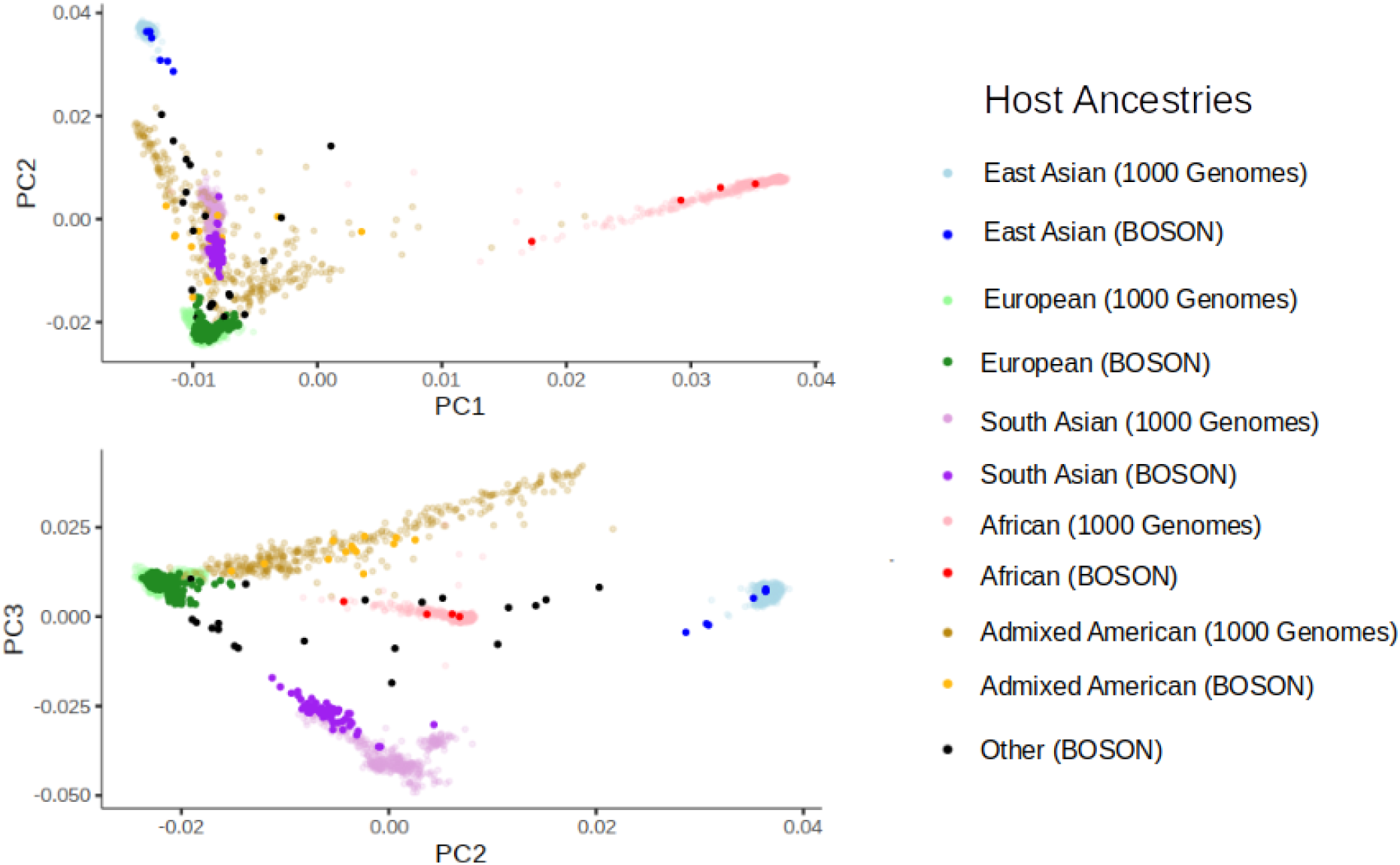
BOSON cohort host genetic principal components projected onto the first three principal components of the 1000 Genomes project. The points in the plots are color coded by ethnicities of individuals in the 1000 Genomes project and host ancestries after adjusting the self-reported ethnicities in the BOSON cohort using ethnicity data from the 1000 Genomes project.

Although the patients with HCV-3a in the BOSON cohort were recruited from the United Kingdom (N=211), the USA (N=65), Canada (N=64), Australia (N=113) and New Zealand (N=35) which are mainly of white European ancestry, we observed an enrichment of patients of South Asian ancestry in this cohort, especially in the United Kingdom (19%, 41/211, p-value = 1.4×10^−13^), where South Asians make up 5% of the population).

### The genetic diversity of HCV-3a is higher in individuals of South Asian ancestry

We used the whole viral genomes to estimate a maximum likelihood (ML) tree. As shown in Figure 2A, the tree consists of four major clades each with a distinct composition of host demographics. We found that when we distinguished patients of South Asian ancestry from other patients, almost all of the viruses from hosts of South Asian ancestry irrespective of their sampling countries grouped together in one clade (“South Asia clade”, the ancestry of 88% of hosts is South Asian in this clade). In the other three clades, the majority of the hosts had white European ancestry and very few South Asian hosts were present. In the “UK clade” 62% of hosts were sampled in the UK, in the “North America clade” 58% of hosts were sampled in North America (USA and Canada) and in the “Australia clade”, Australian and New Zealand samples constituted the largest fraction of the clade (34%). Within each major clade we observed geographical structuring of the isolates. For instance, subclades containing mainly isolates from the Australian continent are dispersed across three of the four major clades, indicating a complex phylogeographical history with multiple independent introductions (Figure 2A). The same pattern is true for North American isolates. The South Asian clade is particularly interesting because it is almost exclusively associated with hosts of South Asian ancestry sampled across different continents. Furthermore, we found that the HCV genetic diversity from South Asian hosts in western countries, is higher than hosts with other ethnicities in these countries and similar to South Asian hosts sampled in South Asia (Figure 2D). Additionally, South Asian hosts in western countries are on average younger than hosts with other ethnicities in these countries (Figure 2C). These observations suggest a distinctive epidemiological history for viruses from South Asian hosts, regardless of sampling locations.

**Figure 2:**
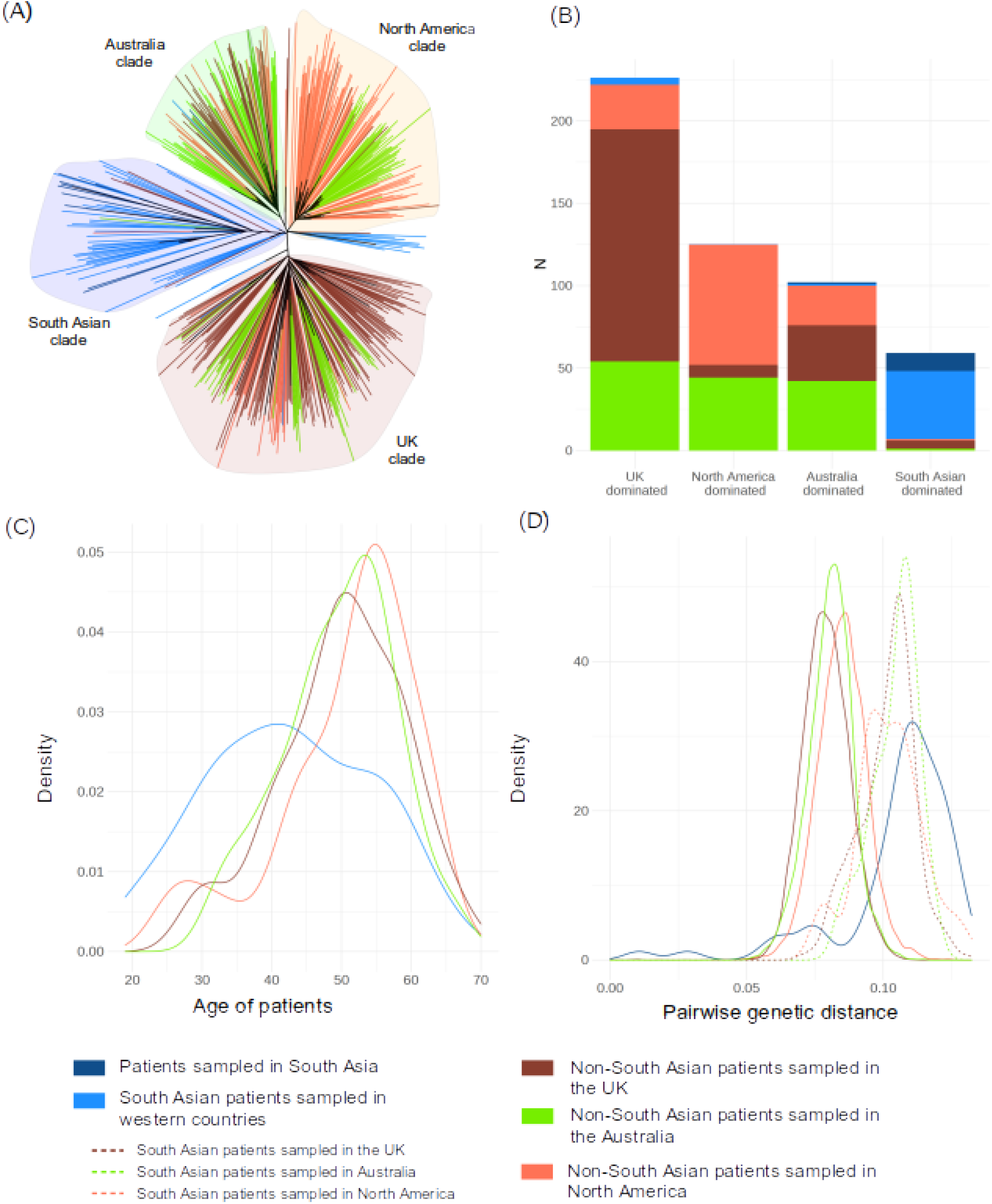
HCV subtype 3a phylogeny and its association with host ancestry and demography. Colours indicate host ancestry and demography. For hosts with South Asian ancestry, their ancestry is indicated irrespective of their sampling location. For the individuals with other ancestries (mainly white European) their sampling location is indicated. (A) Maximum likelihood tree with the tip branches colored by the corresponding host groups. The four major clades associated with distinct demographics are labelled (B) Composition of host ancestry and demography for each of the major clades. (C) Age distribution in different host groups (age data only available for hosts from the Boson Cohort). (D) Distribution of pairwise genetic distance between HCV whole genomes in different host groups.

It has been shown that HCV genotype 3 is endemic in South Asia and it is likely to be the source of the current HCV-3a pandemic (17). Additionally it has been reported that in the UK, South Asian ethnicity is a major risk factor for HCV infection, particularly those that migrated from Pakistan (27–30), and HCV in the South Asian community in the UK has been a public health concern (31). It has been reported that in the UK among non-intravenous drug users, people of South Asian ancestry make up nearly half of all the HCV infections (they constitute ~5% of the population), with South Asian migrants having a 5-fold increased risk over people of South Asian ancestry born in the UK (28). In addition, a history of having surgery, dental/medical treatment, or injection in South Asia were also identified to significantly increase the risk of HCV infection (32). Therefore, we hypothesise that many individuals with South Asian ancestry diagnosed with HCV infection in Western countries are likely to have contracted HCV in South Asia, either due to travel history to the region or living in the region before migration. Under this hypothesis, we would expect hosts that contracted the virus in South Asia to be infected by HCV that are highly diverse and group close to the root of the phylogenetic tree, regardless of their sampling locations. To investigate this, we inferred a time-calibrated ML tree and a Bayesian time-calibrated maximum clade credibility (MCC) tree. The patterns of the ML and

MCC trees showed that the majority of HCVs from individuals with South Asian ancestry living in the Western countries coalesce near the root of the tree and nest the isolates from individuals with other ancestries (Figure 3). This suggests significant discordance between sampling locations and the locations of infection amongst the individuals of South Asian ancestry living in Western countries. Such discordance can potentially confound the inference of HCV-3a epidemiological history. To distinguish between sources of HCV infection in individuals with South Asian ancestry living in Western countries from their sampling location, we reconstructed the ancestral states regarding the host ancestries of HCV by applying structured coalescent analysis to host ancestries (Figure 3). Any HCV-3a isolate from a host of South Asian ancestry sampled in a western country where all its ancestral nodes were also estimated to have South Asian ancestry hosts (in either the ML or the MCC trees) were inferred to have been contracted in South Asia instead of their sampling locations. The locations of infection for other HCV isolates were inferred to be the same as their sampling locations. This resulted in 52 South Asian hosts sampled in western countries identified as having been infected in South Asia (Figure 3) and 5 South Asian hosts sampled in western countries identified as having been infected in their sampling countries.

**Figure 3:**
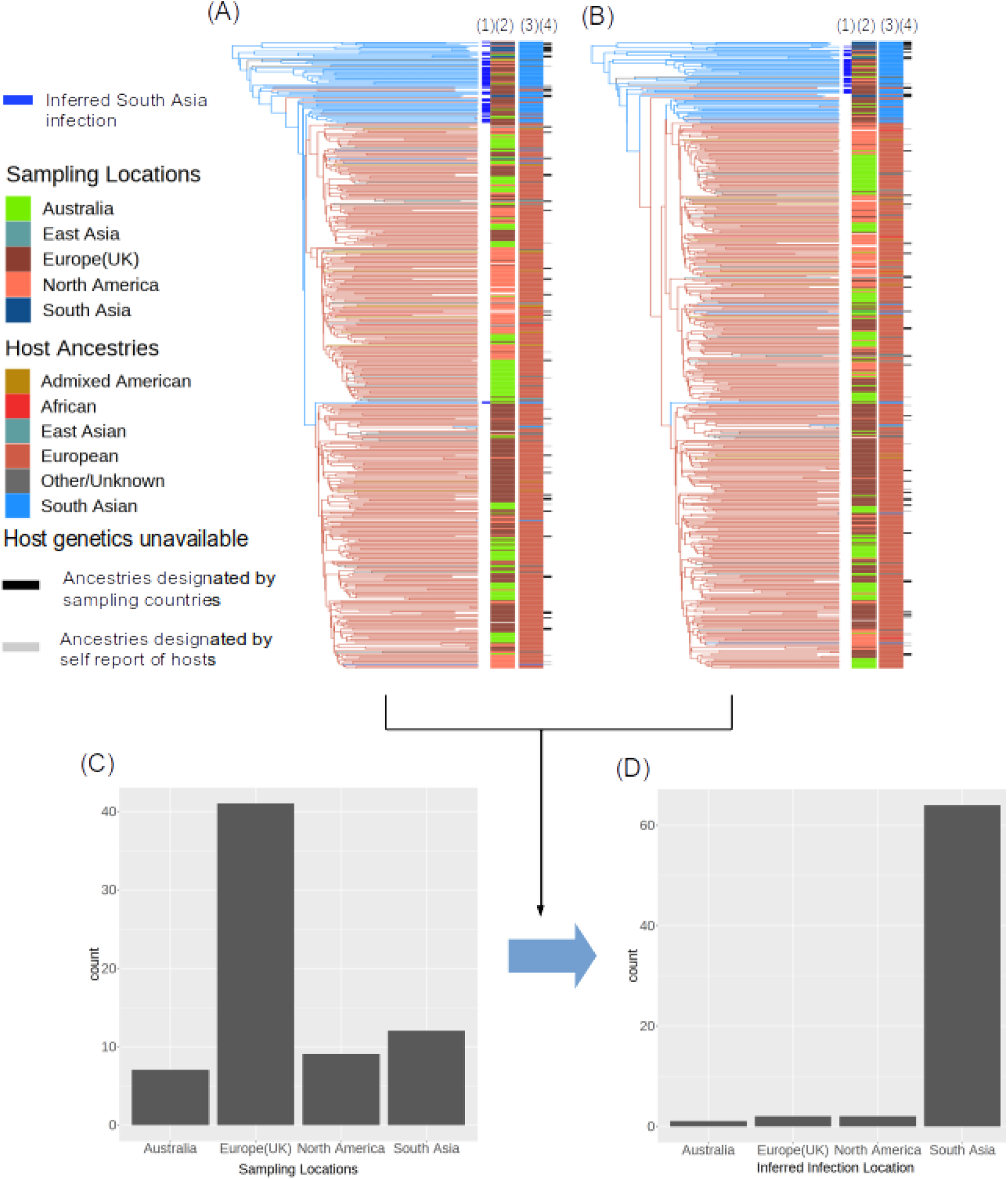
Structured coalescent phylogeographic analysis. Applied to ML (A) and MCC trees (B) to infer the location of infection for South Asian hosts. The branches are colored by the most likely host ancestral states. Lines on the tips of the trees indicate: (1) South Asian hosts sampled in a Western country and inferred to have been infected in South Asia (2) Sampling locations (3) host ancestries and (4) absence of host genetic information. The two bar charts represent the location distribution for (C) sampling locations, and (D) inferred locations of infection amongst hosts of South Asian ancestry.

### Global spread of three distinct clades of HCV-3a

We conducted phylogeographic analysis based on the MCC tree using the inferred locations of infection obtained in the previous section. The inferred time to most recent common ancestor (TMRCA) of the HCV-3a isolates in our study is 1923 (95% HPD, 1905-1938) and it was located in South Asia (geographic posterior probability 0.99) where HCV-3a is likely to have been endemic. Subsequently, three distinct lineages escaped South Asia and spread globally at around the same time. These independent introductions are from South Asia to the UK around 1954 (95% HPD: 1946-1963, node EU in Figure 4), to North America around 1952 (95% HPD:1942-1961, node NA in Figure 4), and to Australia around 1955 (95% HPD: 1945-1963, node AU in Figure 4) respectively. These three introductions form three major lineages approximately corresponding to the three of the major clades illustrated in the ML tree in Figure 2 (UK, North America, and Australia clades). All these introductions occurred during or shortly after the second World War. After these early transmissions out of South Asia, it is estimated that there were frequent transmissions between continents until the late 1960s (Figure 4). Large numbers of independent transmission events to and from Australian continent are observed during this period.

**Figure 4:**
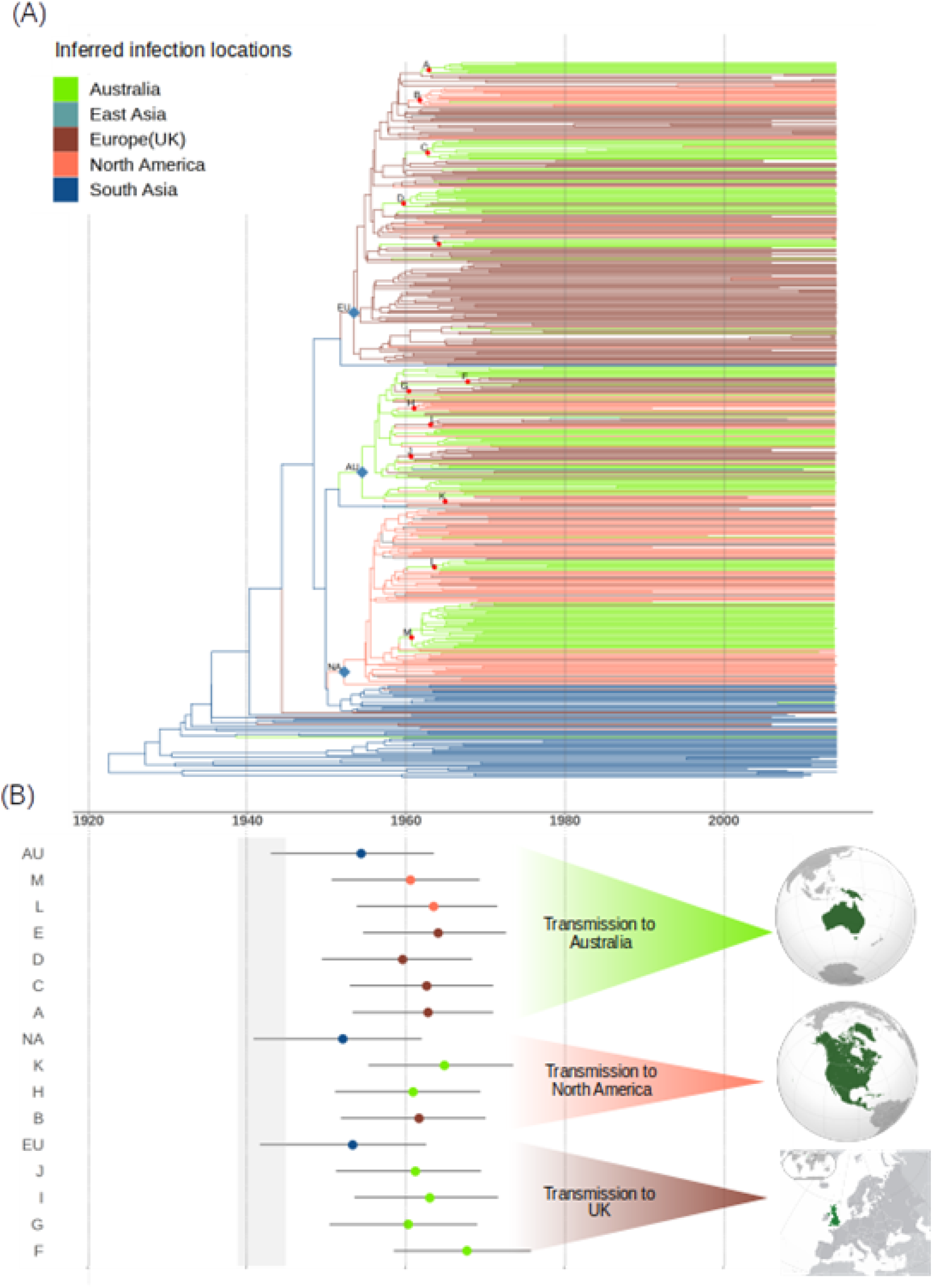
Phylogeographic analysis of HCV epidemiology. (A) MCC tree with branches color coded by the most likely infection location. The three nodes that correspond to the earliest introductions from South Asia to the UK (EU), North America (NA), and Australia or New Zealand (AU) are highlighted in blue. Other significant transmission events between continents are highlighted in red. (B) The 95% HPD of the time of the highlighted nodes. The estimated time points are color coded by the geographical source of transmission. The grey block corresponds to the time period of the second world war (1939–1945).

We investigated the history of the HCV-3a epidemic using a Bayesian skyline plot (Figure 5). Bayesian skyline plot estimates the effective population size of HCV-3a through time. The disease was relatively constant in effective population size through time until the 1960s, when it starts to grow exponentially until the 1980s, after which the growth stops, and the effective population size of the virus becomes constant again.

**Figure 5:**
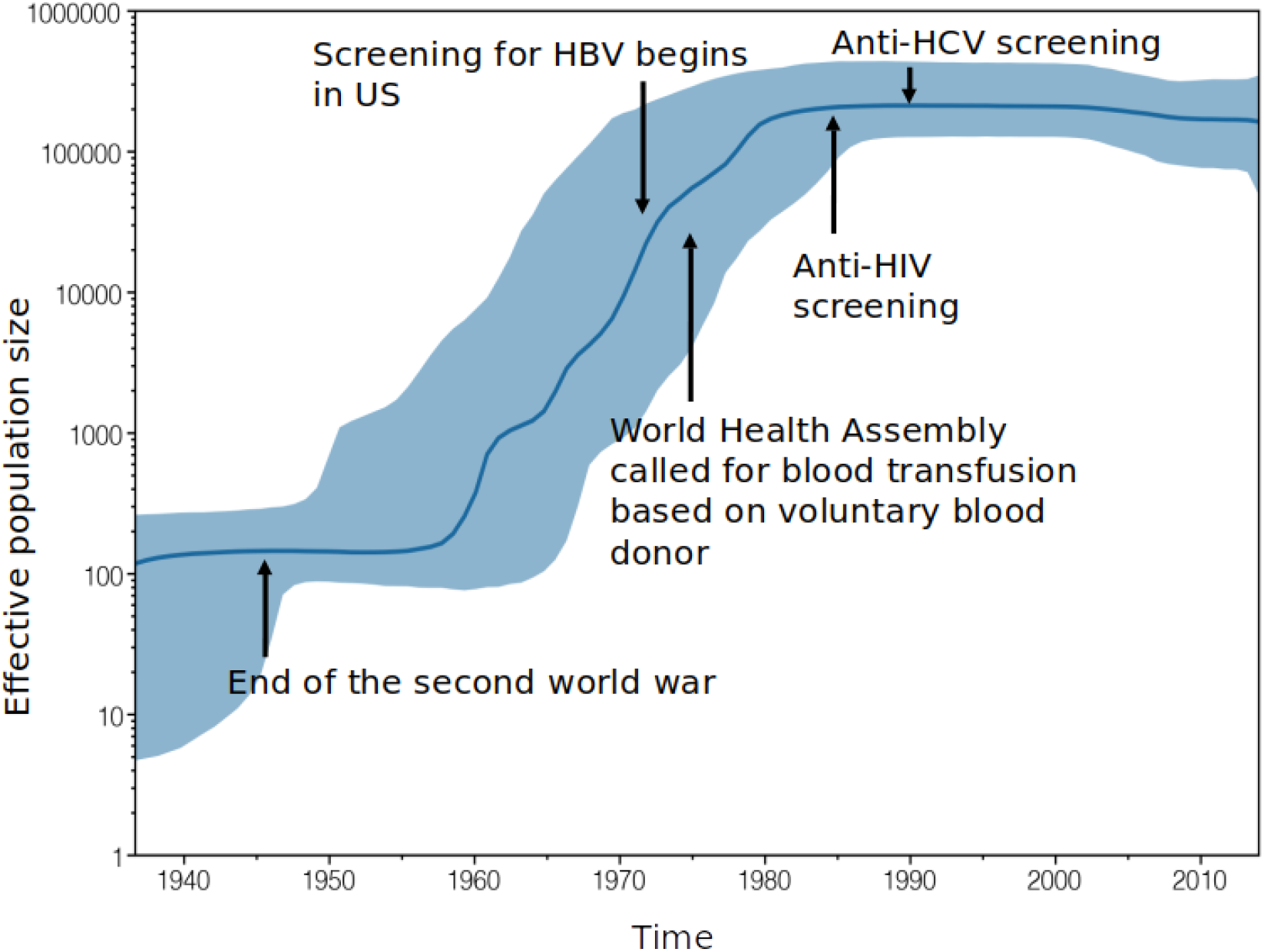
Bayesian skyline plot, showing the epidemic history of HCV subtype 3a. The blue line represents the estimated effective population size through time while the light blue area represents the 95% highest posterior density interval of the estimation. Historical events that might have influenced the growth of HCV-3a population are also shown.

## Discussion

We report the first study which incorporates host genetic information to investigate the genetic epidemiology of a virus (HCV-3a). This additional layer of information allowed us to observe that viruses from South Asian ancestry hosts have a distinct pattern of genetic diversity and that they coalesce near the root of the phylogenetic tree regardless of the sampling countries. The age distribution of South Asian ancestry hosts is also different from hosts with other ancestries and they are over-represented among HCV-3a infected populations in the UK and other countries.

The most likely hypothesis to explain these distinct patterns is that many South Asian patients in western countries are infected in South Asia either before migration or during travels to the region. An alternative explanation would be that these hosts contracted the disease within the South Asian communities in Western countries. However, this scenario is less likely since we would expect the HCV-3a genetic diversity within each of these South Asian communities to be significantly lower than those sampled from South Asia, which is not the case (Figure 2). Also, studies have shown a 5-fold increased risk of HCV infection for South Asians born outside of the UK over UK born South Asians and that the significance of intrafamilial transmission have been controversial (33). Having access to epidemiological data such as the locations of birth and migration history could inform our hypothesis, but in this study, host genetics approximated those epidemiological information which are usually more difficult to obtain. To address the discordance between sampling locations and locations of infections, we conducted a two-step structured coalescent analysis. The first analysis was applied to host ancestry where we leveraged the ancestral states of host ancestries to infer the location of infection amongst South Asian hosts in western countries. The second analysis then used the inferred locations of infection to conduct phylogeographic analysis to rebuild the epidemiological history of HCV-3a. For non-BOSON samples (where we do not have host genetic information), the host ancestry is inferred as the majority ethnic group in their sampling countries. From Figure 3 we can observe that some of these inferred European hosts are very close to the root and inside the South Asian cluster. It is possible that these are actually hosts with South Asian ancestry which were sampled in other countries. Despite this conservative approach, we observed that the coalescence events near the root of the phylogenetic tree are almost exclusively between virus isolates from South Asian ancestry hosts.

We estimated the date for the most recent common ancestor of HCV subtype 3a to be around 1923 (95% HPD: 1905 to 1938), which is the about the same as previous study of HCV-3a by Khan et al. (34), and slightly more recent when compared to some other previous studies of HCV-3a (1899, 95% HPD: 1865-1932 by McNaughton et al. (19); 1914, 95% HPD: 1876-1950 by Choudhary et al. (17)). Another study found the TMRCA of global HCV-3a sequences to be around 300 years ago (23), much earlier than our findings. However, there were four Pakistani sequences that formed a distinct cluster which is distant from other sequences in that study. The rest of the HCV-3a sequences have TMRCA around 1943 (95% HPD: 1885-1958), which is a lot closer to the estimation of our study. The evolutionary rate was estimated to be around 1.694 × 10^−3^ (95% HPD: 1.41 × 10^−3^ to 1.96 × 10^−3^) substitution per site per year, which is close to previous finding by McNaughton et al. (1.65 × 10^−3^) (19), and slightly higher than the estimation by Zehender et al.(1.3 × 10^−3^) (23). Phylogeographic analysis identified the root of the phylogenetic tree to be in South Asia (posterior geographic probability = 0.99). The finding of a South Asian origin agrees with previous studies on the evolutionary history of HCV genotype 3 (15–17,34,35). We then identified three independent introductions of HCV subtype 3a from South Asia to the UK, North America (USA and Canada), and Australia (and New Zealand) that correspond to the beginning of the global spread of HCV-3a. All of these introductions were estimated to be in the 1950s, shortly after the end of the second world war. During the war soldiers from the commonwealth countries fought side by side. HCV is likely to have been transmitted from South Asian soldiers to other commonwealth soldiers such as British, Australians, Canadians and New Zealanders during this time which were then brought back to their home countries after the war. Additionally, after Britain relinquished its Indian empire in 1947, the British Nationality Act of 1948 allowed free entry for citizens of Commonwealth countries into Britain to rebuild the country after the war. As a result, there was a rapid growth in migration from Pakistan and India, two states created by The Partition of India in 1947, to the UK. This migration route could be an alternative explanation for the introduction of HCV-3a from South Asia to the UK.

From the 1960s a series of complex and frequent transmissions between continents followed, presumably due to the growth in global travel. Many of these transmissions were to and from Australia, which was promoting mass immigration under the “populate or perish” policy with a large number of migrants from the UK. This time period also coincides with the start of the exponential growth of the HCV subtype 3a pandemic. Studies have linked various historic events during this period to the HCV expansion. This includes the rise of intravenous drug use (IDU) observed in Europe (36), which has been linked with the growth of HCV subtype 1a and 1b (18). Since IDU has been shown to be the main transmission route for HCV genotype 3 (37,38), we suspect that these events indeed facilitated the spread of the disease during this period, and could have been intensified by blood transfusion and unsafe therapeutic injections (18). The growth of the pandemic stopped in the 1980s. A similar pattern of growth has been observed in HCV subtypes 1a and 1b (18), where the end in population size growth was attributed to the rise of awareness of post-transfusion hepatitis, which led to better regulations around blood donation and needle sharing. During this time, voluntary blood donation, which has been shown to be safer than paid blood donation due to the difference in the demographics of the donors, was promoted in the United States as well as European countries. Also, while this is before the anti-HCV screening in the early 1990s, screening for HBsAg effectively reduced blood donation from high risk populations (39).

In this study we used the largest HCV subtype 3a whole genome dataset to date and combined it with the host genetic data and reconstructed virus epidemic history and identified its origin in South Asia. We observed three independent introductions from South Asia to the UK, North America and Australian continents. Furthermore, we identified that the global spread of the HCV subtype 3a started soon after the end of the second world war and observed many independent transmission events between the UK, North America and Australia after the initial introductions from South Asia. Improvement in technologies and reduction in cost of high throughput technologies has meant that both host and virus genomic information can be obtained on patients. This additional layer of information can be very informative in studying pathogen epidemic history, especially chronic infections where there is a potential dissociation between the sampling location and the location of infection.

## Materials and Methods

### Materials

507 HCV-3a whole genome sequences were obtained from the BOSON clinical trial study that has been described earlier (24). However, the sequences from the BOSON study were all sampled in a short time span. As a result, these sequences have insufficient temporal signal to conduct a meaningful molecular evolution analysis. To address this, we downloaded previously published HCV sequences in the Los Alamos HCV database (40). The criterion for choosing the sequences to download are as follow:

1. The sequence has to be categorized as HCV genotype 3a.
2. The sequence has to cover the coding region of the genome.
3. The sequence has to have sampling date information available.
4. When there are multiple sequences sampled from the same patient at different time points, we only include the earliest sample.
5. When there are multiple sequences sampled from the same patient at the same time point, we randomly select one sequence to include in our dataset.

This resulted in 48 sequences that we then included into our dataset.

### Quality control

The 555 whole genome sequences (from the BOSON cohort and the downloaded sequences) were aligned using MAFFT (41). We then used FastTree (v2.1.10)(42) to build a maximum likelihood (ML) tree. This tree was then used in Tempest (43) to explore the signal of the molecular clock by conducting regression on the root-to-tip genetic distance against sampling time. This allowed us to identify and remove 9 outliers whose root-to-tip genetic distance deviates from the expectation of the linear model by more than 0.025 substitutions per site. The remaining sequences were then realigned using MAFFT (41). Hypervariable regions (HVR)1 and 2 in the sequences were removed from the aligned sequences to avoid downstream analysis from being confounded. We then investigated the aligned sequences by looking at their ML tree as constructed by FastTree (42). We observed that several pairs of sequences on the tips of the tree were very closely related (Sup. Fig. 3). Upon further investigation, we found that all these pairs of samples were from the same countries. Furthermore, all these samples were downloaded from Los Almos database, and many of them had little description available in GenBank and were submitted by the same authors (Sup. Table 1). This raised the concern that these pairs of closely related samples were obtained from the same patients. To be conservative, we randomly discarded one sample in each of these closely related sample pairs. As a result, 6 more sequences are removed from our dataset.

### Host ancestry designation

We downloaded unimputed 1000 Genomes Project phase 3 genetic data (26). Using Plink (v1.9) (44), the SNPs in 1000 Genomes Project were pruned by linkage disequilibrium (LD) on all autosomes and the same set of SNPs were also pruned from the host genetic data from the BOSON cohort. Principal components (PCs) were then estimated for both 1000 Genomes Projects and BOSON respectively using Plink (v1.9) (44). We then projected the PCs of the BOSON cohort on the PCs of 1000 Genomes Project. By comparing the self-reported ancestries in the BOSON cohort to the individuals of the 1000 Genome Project, we found that South and East Asian were both being reported as “Asian”, and some patients that self-reported as “White” were distant from the European ancestry group of the 1000 genome samples in the PC coordinates. To address this, we trained a random forest classifier as implemented by R package randomForest (45) on top 20 PCs from the 1000 Genome Project. We applied this random forest on the PCA data of the BOSON cohort to assign ancestries if the random forest probability was >60%. For individuals where the random forest probabilities were <60%, the “Other/Unknown” ancestry was assigned. For BOSON hosts where genetic data is unavailable (N=19), we used the following scheme for host ancestries designation: We inferred hosts with self-reported ethnicity of “Asian” (N=3) to be of South Asian ancestries as most self-reporting Asians in BOSON are of South Asian ancestries (84%) according to the result of PCA; We inferred hosts self-reporting as “White” to be of European ancestries (N=15); We designated hosts with other self-reported ethnicity as “Other/Unknown” ancestry. For sequences that are downloaded from Los Alamos HCV database, the host genetic information is not available to us. We therefore assigned their host ancestries as the major ancestry group of the countries they were sampled from.

### Phylogenetic analysis

The molecular clock, demographic change, and Bayesian coalescent-based phylogeny of HCV were estimated using BEAST (v1.10.4)(25), which infers rooted and time-scaled phylogenies using clock models via a Markov chain Monte Carlo (MCMC) algorithm. Because of the large sample size in our dataset, we took a few steps to ensure that the MCMC computation could be finished within a reasonable time frame. We first used FastTree (42) to build a ML tree, then used tempest (43) to root the tree. Tempest searches for the most likely root by finding the point in the tree that maximizes the likelihood of the tree given the sampling dates of the tips. The rooted maximum likelihood tree was then used with the R package TreeDater (46) to estimate a time-scaled tree. The resulting tree was then used as the starting point for the MCMC analysis conducted by BEAST. We used the HKY substitution model with a Gamma rate heterogeneity model and the uncorrelated relaxed clock model to model the process of sequence evolution, and the constant population size model for the coalescent process. The “new tree operator mix” option that is available in BEAUti (47), which is a graphical user interface application for generating XML files used by BEAST, was chosen for specifying the moves conducted by MCMC. Finally, the chain length of the MCMC was set at 100 million. MCMC analysis by BEAST was then run twice and Tracer(v1.7.1)(48) was used to visually inspect the result of the MCMC analysis to ensure good mixing. Beside Bayesian coalescent-based phylogeny, we also used a maximum likelihood approach to estimate the phylogeny of HCV. RAxML (v8.2.12)(49) was used to estimate a maximum likelihood tree which was then rooted with tempest (43). The tree was then dated by treeDater. The BEAST2 (50) package MASCOT(51) was used for phylogeographical analysis, where the phylogeny was fixed, population sizes and migration rates between populations were set to be uniform, and the MCMC chain length was set to 10 million steps for each MASCOT run.

