## Supplementary Material for "Global spread and evolutionary history of HCV subtype 3a"

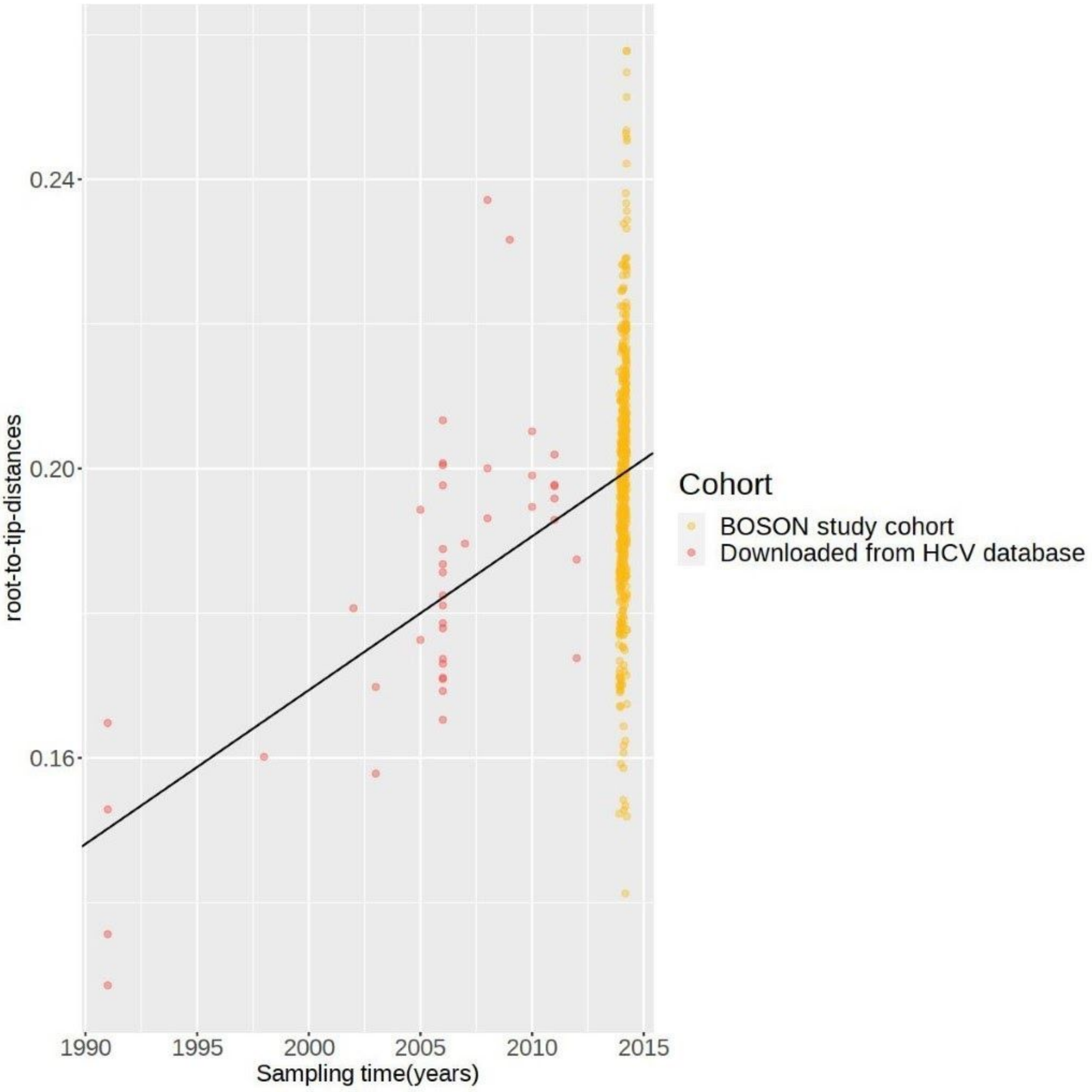


Supplementary Figure 1: Molecular clock analysis and the temporal distribution of HCV-3a samples. A maximum likelihood tree generated from whole genomes was used to estimate root to tip distance for each isolate which was then plotted against the sampling date and a regression line was fitted to the data.

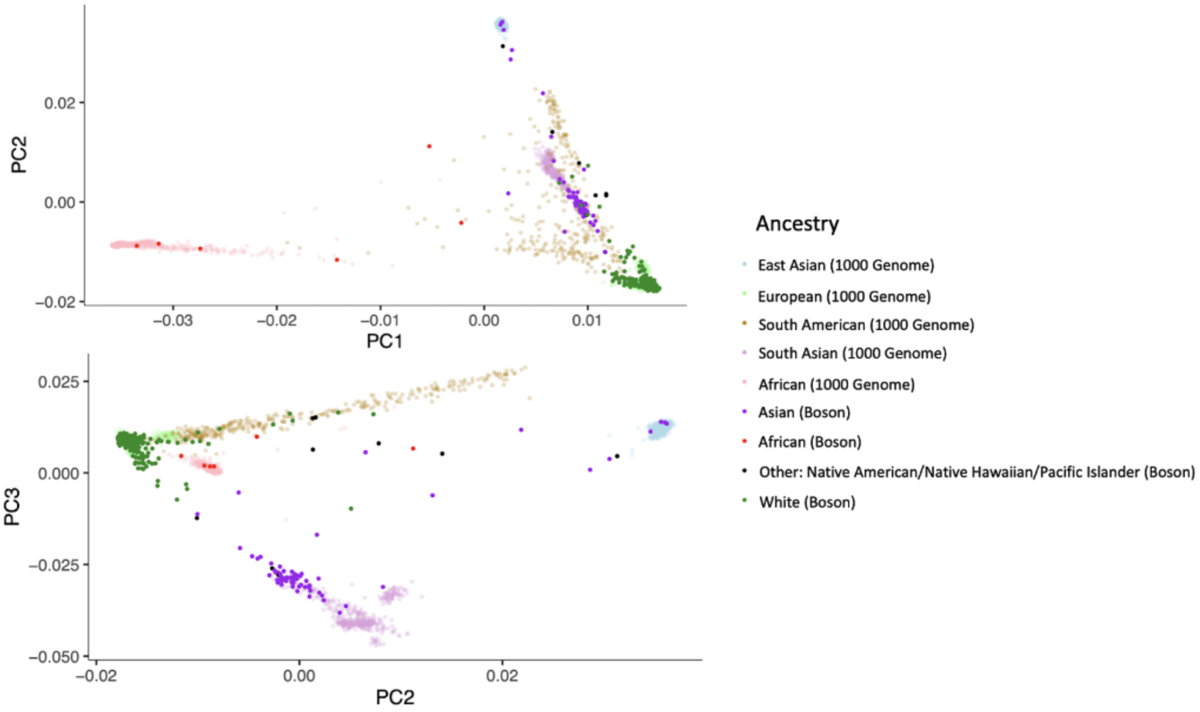


Supplementary Figure 2: Scatter plots of the BOSON cohort host genotypes projected onto the first three PCs calculated from the 1000 Genomes Project. The points in the plots are color coded by ethnicities of individuals in the 1000 Genomes project and BOSON self-reported ethnicities.

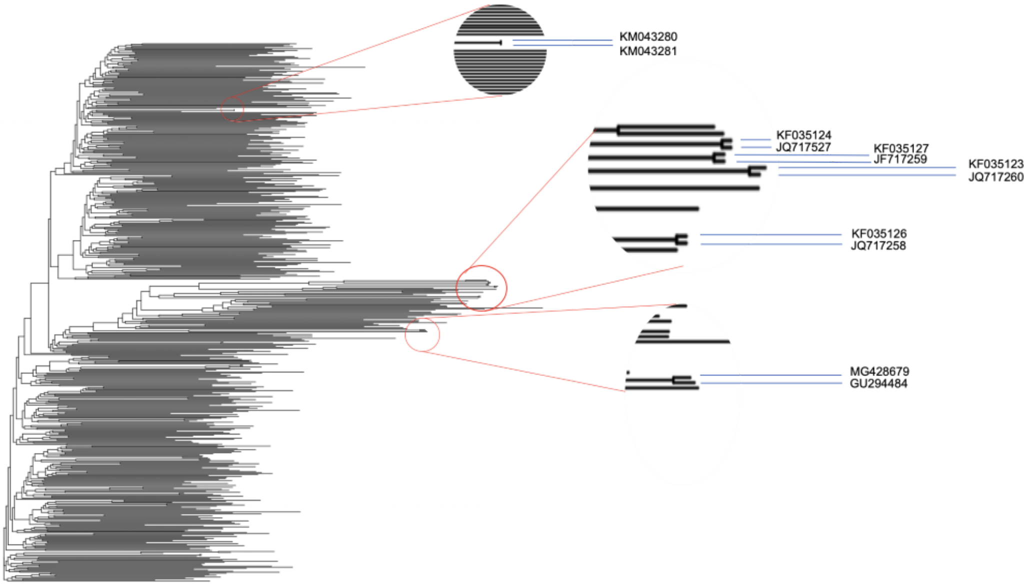


Supplementary Figure 3: The samples that we identified as potential duplicates, illustrated in an unrooted ML tree.

| Duplicated pair | Authors | Sampling location |
|---|---|---|
| MG428679 | Ahmad,S., Ali,I. and Ahmad,S. | Pakistan |
| GU294484 | Rehman,I., Butt,S., Idrees,M., Rafique,S., Akbar,H., Zubair,M., Awan,Z., Manzoor,S., H usain,A., Akram,M., Khubaib,B. and Aftab,M. | Pakistan |
| KM043280 | Stoddard,M.B., Li,H. and Shaw,G.M. | USA |
| KM043281 | Stoddard,M.B., Li,H. and Shaw,G.M. | USA |
| KF035126 | Choudhary,M.C., Natarajan,V., Mishra,G., Tripathi,R., Gupta,E. | India |
| JQ717258 | Choudhary,M.C., Mishra,G., Tripathi,R., Gupta,E., Singh,T. | India |
| JQ717260 | Choudhary,M.C., Natarajan,V., Pandey,P., Gupta,E., Sharma,S., Tripathi,R., Kumar,M. S., Kazim,S.N. and Sarin,S.K. | India |
| KF035123 | Choudhary,M.C., Natarajan,V., Mishra,G., Tripathi,R., Gupta,E., Singh,T., Trehanpati,N., Kazim,S.N., Kumar,M.S. and Sarin,S.K. | India |
| JQ717257 | Choudhary,M.C., Natarajan,V., Pandey,P., Gupta,E., Sharma,S., Tripathi,R., Kumar,M. S., Kazim,S.N. and Sarin,S.K.. | India |
| KF035124 | Choudhary,M.C., Natarajan,V., Mishra,G., Tripathi,R., Gupta,E., Singh,T., Trehanpati,N., Kazim,S.N., Kumar,M.S. and Sarin,S.K | India |
| KF035127 | Choudhary,M.C., Natarajan,V., Mishra,G., Tripathi,R., Gupta,E., Singh,T., Trehanpati,N. , Kazim,S.N., Kumar,M.S. and Sarin,S.K. | India |
| JQ717259 | Choudhary,M.C., Natarajan,V., Pandey,P., Gupta,E., Sharma,S., Tripathi,R., Kumar,M.S., Kazim,S.N. and Sarin,S.K. | India |
